# An evolutionary model identifies the main selective pressures for the evolution of genome-replication profiles

**DOI:** 10.1101/2020.08.20.259416

**Authors:** Rossana Droghetti, Nicolas Agier, Gilles Fischer, Marco Gherardi, Marco Cosentino Lagomarsino

## Abstract

Recent results comparing the temporal program of genome replication of yeast species belonging to the *Lachancea* clade support the scenario that the evolution of replication timing program could be mainly driven by correlated acquisition and loss events of active replication origins. Using these results as a benchmark, we develop an evolutionary model defined as birth-death process for replication origins, and use it to identify the selective pressures that shape the replication timing profiles. Comparing different evolutionary models with data, we find that replication origin birth and death events are mainly driven by two evolutionary pressures, the first imposes that events leading to higher double-stall probability of replication forks are penalized, while the second makes less efficient origins more prone to evolutionary loss. This analysis provides an empirically grounded predictive framework for quantitative evolutionary studies of the replication timing program.

## I. INTRODUCTION

Eukaryotes, from yeast to mammals, rely on pre-defined “replication origins” along the genome to initiate replication [1–4], but we still ignore most of the evolutionary principles shaping the biological properties of these objects. Binding by initiation complexes defines origins as discrete chromosomal loci, which are characterized by multiple layers of genomic properties, including the necessary presence of autonomously replicating sequences, nucleosome depletion, and absence of transcription[5, 6]. Initiation at origins is stochastic, so that different cells of the same population undergoing genome replication in S-phase will typically initiate replication from different origins [7, 8].

Initiation from a single origin can be described by intrinsic rates and/or licensing events [9]. Indeed, the genome-wide replication kinetics of a population of cells can be accessed experimentally by different techniques [9–11]. Recent techniques also allow to measure replication progression at the single-cell level [12, 13]. The estimation of key origin parameters from data requires minimal mathematical models describing stochastic origin initiation and fork progression [10, 14–16]. Typically, one can extract from the data origin positions, as well as estimated origin-intrinsic characteristic firing times or rates. Knowledge of origins positions and rates makes it possible to estimate the “efficiency” of an origin, i.e. its probability of actively firing during S-phase, rather than being passively replicated.

Over evolution, a genome modifies its replication timing profile by “reprogramming” origin positions and rates in order to maximize fitness, under the constraints of the possible changes of these parameters that are physically and biologically accessible. Little is known about this process, and finding basic rules that drive origin evolution is our main focus here [17]. The main recognized constraint determining selective pressure is origin stalling [18–20]. If two replication forks stall with no origins in between them, it is generally agreed that replication cannot be rescued, and the event leads to cell death. A pioneering study by Newman and coworkers [18] used a combination of data analysis and mathematical models to understand the role of lethal double stall events on origin placement. They found that the fork per-base stall probability affects the distance between neighbor origins, and the optimal distance distribution tends to a regular spacing, which is confirmed by experimental data. Thus, origin placement is far from uniform, and appears to be the result of an effective “repulsion” process.

Due to the streamlined genome and the experimental accessibility, yeasts are interesting systems to study experimentally the evolution of replication programs. However, at the level of the *Saccharomyces* genus, the replication program is highly conserved [21]. Hence, until recently, no experimental account of the evolution of the replication program was available. Our collaboration has recently produced data of this kind [22], by comparing replication dynamics and origin usage of 10 distant *Lachancea* yeast species. This study highlights the dominance of origin birth-death events (rather than e.g. chromosomal rearrangements) as main evolutionary drive of the replication program changes, and characterizes the main principles underlying origin birth-death moves. Briefly, origin gains tend to associate with “weaker” (low efficiency) origins, and, while overall old origins tend to be lost more frequently than recent ones, old conserved origins tend to be “stronger” (high efficiency). Additionally, proximity to efficient origins correlates with origin loss events. These findings open the question of capturing the relevant selective pressures acting on replication profiles in the framework of the empirical birth-death evolutionary dynamics, for which the data set [22] provides an empirical testing ground.

Here, we define a minimal evolutionary birth-death model for replication program evolution encompass all the empirical observations made by Agier and coworkers [22], and we use it to identify the main selective pressures and evolutionary trade-offs at play.

## II. RESULTS

### Experimental data motivate an evolutionary model for origins turnover

This section presents a reanalysis of the experimental data from ref. [22]. We summarize the main results of that study, and present additional considerations on the same data, which motivate the evolutionary model framework used in the following.

Supplementary Fig. S1 recapitulates the *Lachancea* clade phylogenetic tree used in the analysis. The evolution of the temporal program of genome replication can be quantified by the divergence of the replication timing profiles across different species. Agier and coworkers found that timing profiles diverge gradually with increasing evolutionary divergence between species [22]. In principle, such divergence could be attributed to changes in the number, placement, and biological properties of all origins. However, a careful analysis of correlations (comparing the timing profiles and the activity of orthologous origins) shows that the main driver of program differentiation across species is the acquisition and loss of active replication origins. Specifically, the number of conserved origins decreases with increasing phylogenetic distance between species, following the same trend as the conservation of the timing profiles. This trend is the same in regions that are close to or away from breakpoints, pointing to a secondary role of genome rearrangements. In addition, the authors of ref. [22] show that the differences in the mere number of origins and the median difference in origin replication timing between pairs of species are nearly constant with phylogenetic distance, leading to exclude that origin reprogramming (rather than birth-death) plays a primary role in the evolution of the timing program.

Any model for the evolution of the replication program must (i) reproduce the empirical distribution of the inter-origin distances, (ii) reproduce the empirical distribution of the origin efficiencies, and (iii) account for the observed origin turnover dynamics. Previous analyses [18, 22] have shown that origins are far from following a uniform distribution along the genome. Fig. 1A shows that the inter-origin distance distribution robustly shows a uni-modal shape across the ten *Lachancea* species studied in ref. [22]. Specifically, distributions for each species show a marked peak around 35 Kbp. This peak corresponds to a typical inter-origin distance, which is strikingly invariant across all *Lachancea* species. Fig. 1B shows the distribution of the efficiencies, which is defined as the probability to actively fire during the S phase, estimated for each origin in the *Lachancea* clade using Eq. 4 and a fit inferring the firing rates of all origins assuming a standard nucleation-growth model (see Methods and ref [16]). The single-species efficiency distributions show more variability across species than the inter-origin distance distributions, but they are consistent with a common shape and support.

**FIG. 1.**
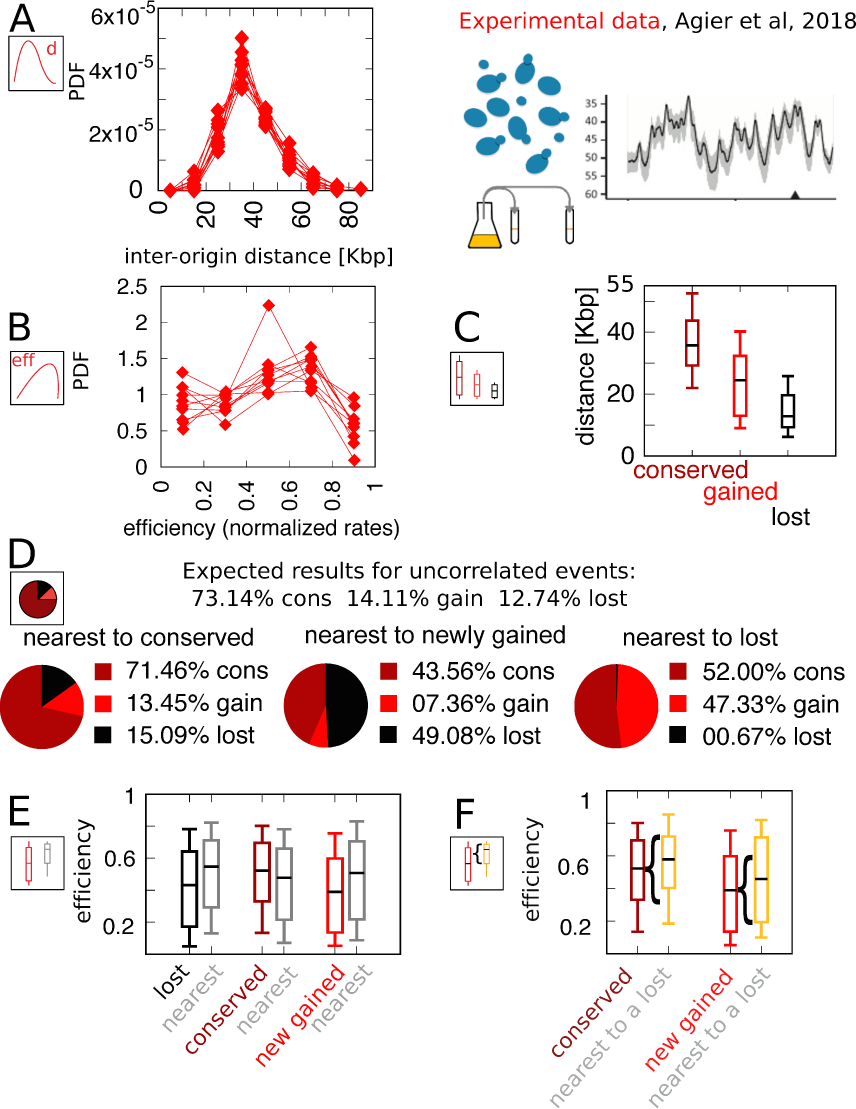
Experimental data motivate an evolutionary model for replication origins turnover. **A:** Distribution of the distance between neighbor origins in ten *Lachancea* species, each histogram refers to a different species (data from ref. [22]), and all the plots show a marked peak around 35 Kbp. **B:** Distribution of the efficiency (calculated from a fit, using Eq. 4) for all origins in ten *Lachancea* yeast species. clade [22]. **C:** From ref. [22], box plot of the distribution of the distance from the nearest origin split by evolutionary events, for conserved (dark red), newly gained (red) and lost origins (black), estimated comparing six sister species of the *Lachancea* clade [22]. **D:** Analysis of the origins that are nearest to conserved, newly gained and lost, compared to the expected result if events were uncorrelated [22]. **E:** Distribution of the efficiency of lost, conserved and newly gained origins (respectively in black, dark red and red) and their neighbors (grey). Note that the efficiency of lost origins is lower than average, while the efficiency of origins flanking a lost origin is higher. **F:** Box plot of efficiency of all conserved and newly gained origins compared to those flanking a lost origin, which tend to be more efficient. Box plots show the median (bar), 25-75 (box), and 10-90 (whiskers) percentiles. The data in panel C, D, E and F refers to the six sister species of the *Lachancea* tree.

As mentioned above, a key result of Agier and coworkers is the insight that the evolution of the replication program is mainly shaped by the birth-death process of replication origins. Fig. 1C-F recapitulate the main quantitative results that characterize this process. Note that the analyses in Fig. 1C-F have been performed on the six sister species of *Lachancea* clade. The other species pairs are too distant to perform a reliable identification of conserved, newly gained and lost origins [22].

Fig. 1C shows a box plot of the distance from the nearest origin for all the conserved (grey), newly gained (green) and lost (red) origins. Lost replication origins tend to be closer to their neighbors, much more so than newly gained or conserved origins. This observation reveals that the distance of an origin from its nearest neighbor is correlated to the loss rate of the same origin over evolution. This is an essential feature that any evolutionary model of this process must take into account [18, 22]. More in detail, Fig. 1D further quantifies the correlation between gain and loss events of neighboring origins, by comparing the fraction of observed events of loss, gain, or conservation, given the state of the nearest origin (conserved, lost, or gained). The distribution of event types for origins that are nearest neighbors of a newly gained origin deviates significantly from the null expectation of random uncorrelated events (i.e., in a simple scenario where the fractions of conserved, newly gained, and lost origins are fixed to the empirical values, and birth and death events of neighboring origins are independent). The same non-null behavior is observed for origins that are nearest to a lost origin, with the roles of gain and loss events exchanged. In summary, successive birth/death or death/birth events happen more frequently in the same genomic location than expected by chance. Beyond such a spatial correlation along the chromosomal coordinate, the analysis illustrates that birth and death events are correlated in time as well (in fact, the analyzed evolutionary events took place in the terminal branches of the phylogenetic tree, and thus they must have been close in term of evolutionary time).

Finally, Fig. 1E and 1F show that origins lying near loci where origins were recently lost are typically in the high-efficiency range of the distribution, and that lost origins tend to be less efficient than conserved origins. Fig. 1E compares the distribution of the efficiency of lost, conserved and newly gained origins with the distribution of efficiency of the nearest origins. The efficiency of origins neighboring a loss event is higher than average, while the efficiency of lost origins is lower than average. These results clearly support the influence of origin efficiency on origin death events. This is confirmed by Fig. 1F, which shows the distribution of efficiency of all conserved and newly gained origins. For both classes, considering only those origins that are nearest neighbors to a recently lost origin yields an increase in the efficiency.

Different mechanisms could lead to the correlations described above. Overall, it is clear that origin strength is somehow “coupled” to birth-death events. For example, conserved origins may become more efficient after the loss of neighbor origins, or the birth of new highly efficient origins could facilitate the loss of neighbors, or losing an origin could expedite the acquisition of a new origin nearby. Overall, these results reveal that the origin birth-death process is following some specific “rules” that involve both inter-origin distances and origin efficiency.

Note that the results of Fig. 1E might appear to be incompatible with Fig. 1D, but they are not. Fig. 1E shows that the efficiency of newly gained origins is lower than average, and Fig. 1D shows that the majority of origins that are nearest to a locus with a recent loss event are newly gained. The apparent contradiction arises from Fig. 1E, which shows that the average efficiency of origins close to a lost one is higher than average. This inconsistency is resolved by the analysis shown in Fig. 1F, which shows that origins appearing close to recently lost ones are among the most efficient.

### A birth-death model including replication forks double stall as the main fitness contribution recapitulates the main features of replication origin turnover

The joint stalling of two replication forks in the same inter-origin region along the genome is a well-characterized fatal event that may occur during S-phase. The frequency of this event in a clonal population clearly affects fitness. A previous modeling study [18] focusing on yeast demonstrated that, in order to minimize the probability of a double stall anywhere along the chromosome, origins must be placed in the most ordered spatial configuration, namely all the consecutive origins must be equidistant from each other. However, the previous study did not incorporate this principle into an evolutionary dynamics of origin turnover. Thus, the important question arises of whether how the selective pressure to avoid double stalls is related to origin gain and loss. To address this question, we defined a birth-death model, rooted in the experimental observations discussed in the previous section. This “double-stall aversion model”, described in detail below, biases the turnover of replication origins in such a way that events (in particular birth events) leading to a decreasing double-stall probability are promoted, because they increase the fitness of the cell.

In the double-stall aversion model, the extent to which the acquisition of a new origin changes the probability of a double stall 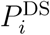 depends on the length *l*_*i*_ of the inter-origin region where the event occurred. This probability is therefore coordinate-dependent, and can be derived by a procedure similar to the one carried out in [18] (see more details in methods),

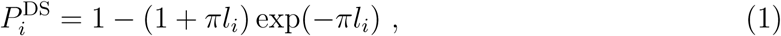

where *l*_*i*_ is the length of the genome region between the (*i* + 1)-th and the *i*-th origin and *π* is the mean per-nucleotide fork stall rate; we use the value from ref. [18], *π* = 5 × 10^−8^ per nucleotide. Note that the double stall probability is completely independent from the origin firing rates and efficiency, and depends only on the distance between the origins.

In our simulations of the model (see methods for a more detailed explanation), the genome was represented as a vector of origins, identified by the position and the firing rate. The model is a discrete-time Markov chain, and for the double-stall aversion variant the chain is specified by the following update rules,

- In each inter-origin region, the origin birth rate is biased by the value of the double-stall probability in that region. Specifically, the origin birth rate (per unit time) in the region *i*, of length *l*_*i*_ between the *i*-th and (*i* + 1)-th origin is given by

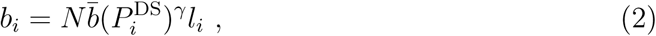

where 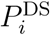 is the (constant) double stall probability density in region *l*_*i*_ (Eq.1), 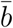 is the birth rate (per Mbp and per unit time) extracted from experimental data (see methods), and *γ* is a positive real parameter that controls the strength of the bias. *N* is a normalization factor added to match the empirical birth rate 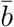. Newborn origins are placed in the middle of the inter-origin region *l*_*i*_.
- Death (i.e., loss of origins) is unbiased, and occurs at random origins with rate 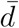 (estimated from experimental data, see methods), regardless of their efficiency or their neighbor’s efficiency.

SI Fig. S5 shows the simulation results of the model with best-fitting parameter values (see methods and Fig. S5 for other parameter values). SI Fig. S5A and B, show that the double-stall aversion model reproduces the two main “structural” features of the replication program, namely the inter-origin distance distribution and the origin efficiency distribution. Additionally, SI Fig. S5C and D show that the same model reproduces the observed correlations between the inter-origin distance and origin birth-death events, as well as the correlation between birth-death events and nature of the neighbor origins observed in the data (conserved, newly gained, or lost).

### The double stall hypothesis alone fails to capture correlations of origin turnover with efficiency

In spite of the good performance of the double-stall aversion model in explaining the empirical marginal distributions, we find that it fails to reproduce the observed correlations between the efficiency of an origin and the recent history of the nearest ones. SI Fig. S5E shows very faint variations in efficiency of origins that are nearest neighbors to origins of different evolutionary fate. In particular, the observed huge divergence in efficiency between lost origins and their neighbors is absent in the model simulations. Note that Fig. 3 and SI Fig. S5F show that in the double-stall aversion model origins nearest to a loss event are slightly more efficient than average. This trend is due to the fact that after an origin is lost, its neighbours are subject to lower interference, and automatically become more efficient. However, Fig. 3 shows that this null trend is too weak to explain the experimental data. These considerations indicate that a model without a direct mechanism linking the efficiency of an origin to the birth-death events of its neighbors cannot reproduce the data.

**FIG. 2.**
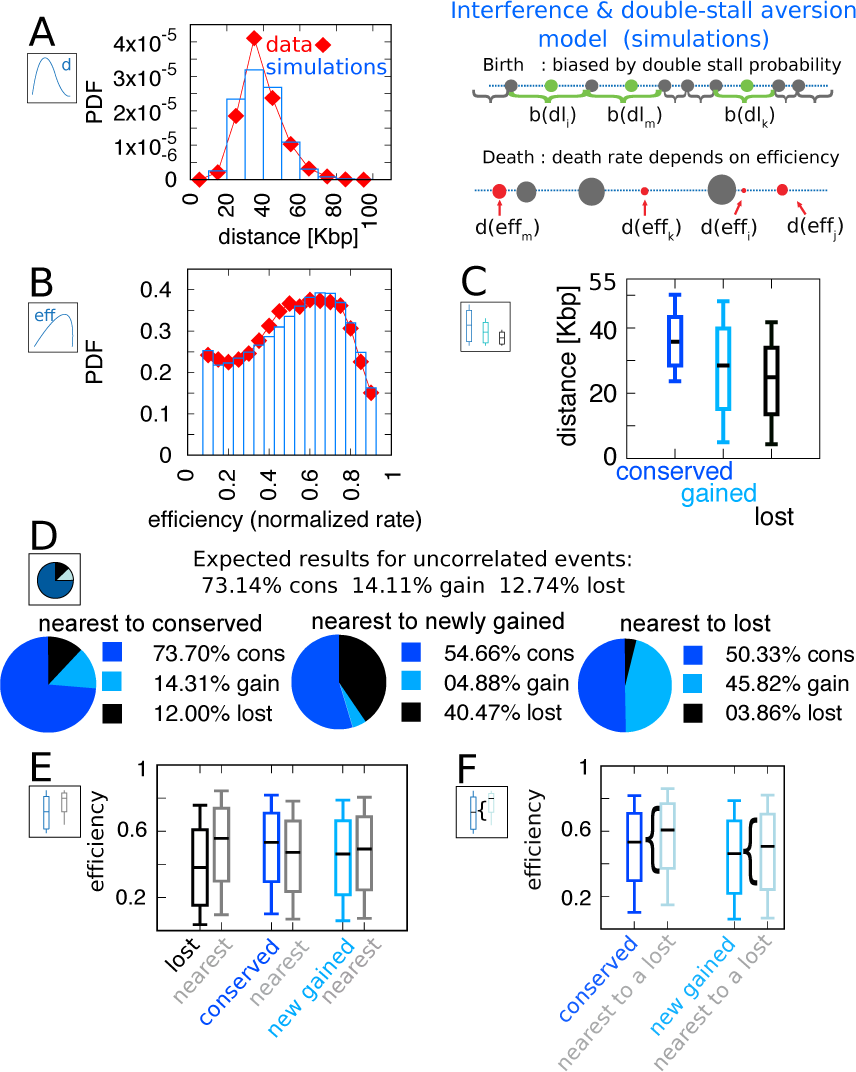
A model where both fork stalling and interference affect fitness explains the correlations between origins evolutionary events. Result of the best-fitting simulation of the joint model compared with empirical data. **A:** Inter-origin distance distribution in simulated species (blue bars) compared to the empirical distribution for the ten *Lachancea* species (red diamonds). **B:** Origin efficiency distribution in simulated (blue bars) *vs* empirical species (red diamonds). The agreement between simulation and experimental data shows that this joint evolutionary model reproduces the typical features of a yeast replication program. **C:** Box plot of the distance from the nearest origin split by evolutionary events, i.e. for conserved (dark blue), newly gained (blue) and lost origins (black), for simulated species. **D:** Fraction of origins that are nearest to conserved, newly gained and lost, for simulated species, compared to the expected result for uncorrelated events. **E:** Box plot of efficiency of lost, conserved and newly gained origins (respectively in black, dark blue and blue) and their neighbors (grey), in simulated species. **F:** The efficiency of all conserved and newly gained origins compared to the ones flanking a lost origin. Box plots show the median (bar), 25-75 (box), and 10-90 (whiskers) percentiles. Panels D and F show that the model correctly reproduces the correlation between origin birth-death events over evolution and efficiency of the nearest origin. Simulation parameters (see methods): *γ* = 2.3, *β* = 1.9, overall birth and death rate 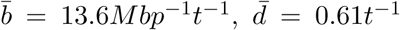 and rate of origin firing-rate reshuffling *R* = 0.92*t*^−1^, where *t* is measured by protein-sequence divergence.

**FIG. 3.**
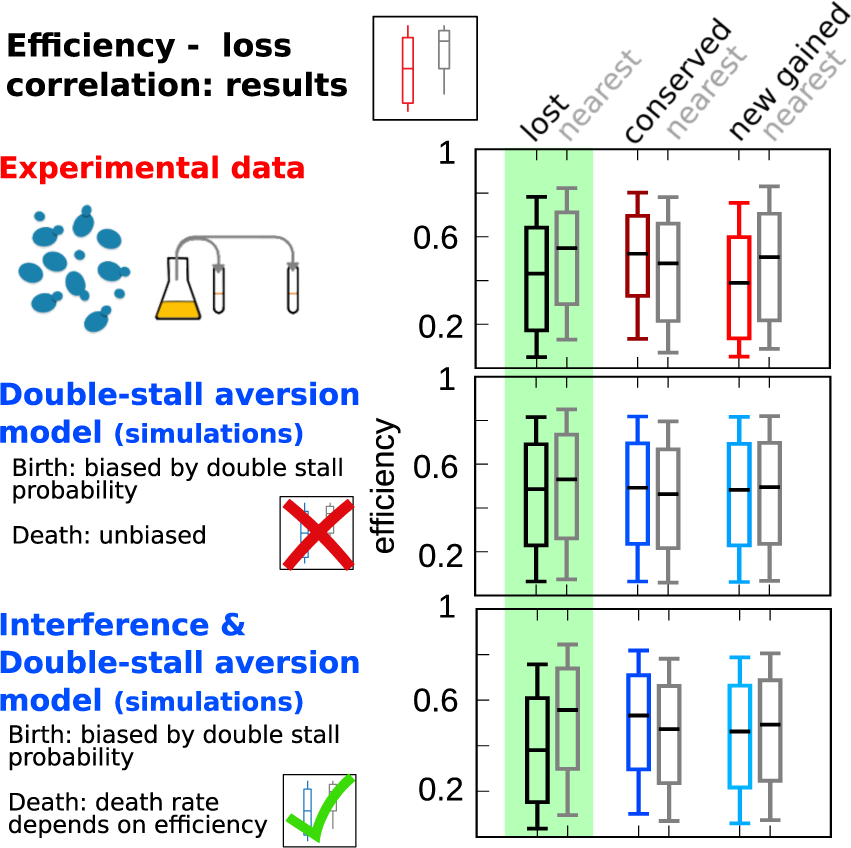
Comparison of model predictions for the correlations of origin birth-death events. The plots compare efficiency distributions of the best-fitting simulation of the two different models (bottom and central panels) with experimental data (top panel). Comparison of the box plot of efficiency of lost, conserved and newly gained origins (red for the data, blue for the models) shows better agreement of the joint efficiency/double-stall aversion model (bottom panel) with the experimental data. Hence, the joint model reproduces well the correlation between evolutionary birth-death events of origins and efficiency of the nearest origin, while the double-stall aversion model fails. Box plots show the median (bar), 25-75 (box), and 10-90 (whiskers) percentiles. Simulation parameters for the joint model (see methods): *γ* = 2.3, *β* = 1.9, and for the double-stall aversion one: *γ* = 2.4. General parameters: overall birth and death rate 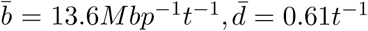 and rate of origin firing-rate reshuffling *R* = 0.92*t*^−1^, where *t* is measured by protein-sequence divergence.

### Double-stall aversion and interference between proximate origins explain the correlated evolution of origin presence and efficiency

Based on the above considerations, we defined a joint model that takes into account both the evolutionary pressure given by the double-stall probability and the direct effect of origin efficiency on birth-death events.

Specifically, this model is defined as follows.

- The birth process is the same as in the double-stall aversion model described above: the birth rate is biased by the double-stall probability in each inter-origin region [eq. (1)], and newborn origin are placed in the middle of the region.
- Death of an origin is biased by its efficiency: less efficient origins are more easily lost. Specifically, the death rate (per unit time) for the *i*-th origin is

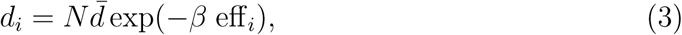

where eff_*i*_ is the efficiency of the *i*-th origin, Eq. (4), 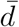 is the mean death rate extracted from experimental data (see methods). The positive parameter *β* tunes the interaction strength: the larger *β*, the steeper the dependence of *d*_*i*_ on eff_*i*_. The normalizing factor *N* is chosen so as to match the empirical total death rate.

Fig. 2 gathers plots of the structural features (distribution of inter-origin distances and efficiencies, Fig. 2A-B) and the evolutionary correlations involving efficiency, evolutionary fate, distance to nearest neighbor, and fate of nearest neighbor (Fig. 2C-D-E-F). Overall, the joint model reproduces all the observations considered here regarding the layout of origins and their evolutionary dynamics, indicating that the experimental data can be rationalized by a fitness function that includes both the detrimental effects of non replicated regions and the evolutionary cost of maintaining inefficient replication origins.

In particular, the coupling between the efficiency of an origin and the death rate of its neighbors, through the probability of passive replication, reproduces the empirical correlations shown in Fig. 1. Figure 3 summarizes this crucial point of comparison between the joint efficiency/double-stall aversion model and the pure double-stall aversion case. The three plots compare efficiency distributions of lost, conserved and newly gained origins (red for the data, blue for the models) with those of their neighbors (grey). Comparison of these plots shows that only the joint model reproduces the differences in efficiency of lost origins and their neighbors.

### The joint efficiency / double-stall aversion model correctly predicts origin family divergence

Having established that the joint model is required to reproduce observations on single lineages, we turned to its predictions on observations that require knowledge of the whole phylogenetic tree, such as origin evolutionary families, defined as sets of orthologous origins [22].

We thus set up a simulation of the model on a cladogenetic structure, fixed by the observed structure of the *Lachancea* clade phylogenetic tree (see methods for the simulations details). The output of each run in such simulations are nine different simulated genomes whose lineages are interconnected in the same way as the empirical species, and each branch follows the empirical divergence. The model for the tree can simulate nine species, all the species except for *L. kluyveri*, as this species was used as outgroup for the computation of the length of the tree branches [22]. We have repeated all the analyses on these simulations, and verified that all the previous results hold, SI Fig. S6. We then turned to other independent predictions of the joint model, which could be compared to measurements in ref. [22].

Fig. 4 reports the dynamics of origin families. As reported in ref. [22], origins that belong to larger evolutionary families tend to have a higher efficiency compared to origins in smaller families. This is related to the observation that older origins tend to be more efficient, and to the fact that the size of a family is roughly related to the age of the origins. Note however that there is no deterministic relation between family size and origin age because the relationship between these two is determined by the structure of the phylogenetic tree. Indeed, two families of the same size may have roots in different points of the tree, and thus the origins belonging to them may have very different ages. Thus, the prediction of the relation between origin efficiency and origin-family size is not trivial. Fig. 4A shows the results for the origin efficiency for families of varying size, comparing the experimental data and 100 different runs of the simulation.

**FIG. 4.**
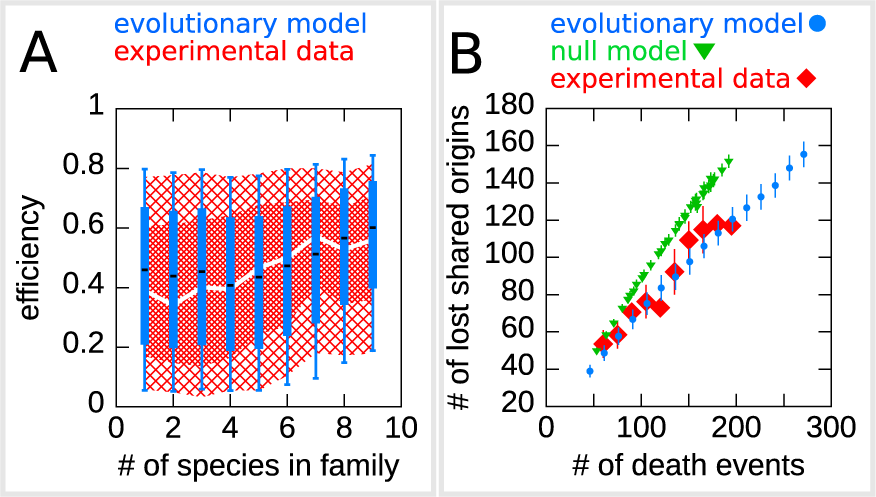
The efficiency/double-stall aversion model predicts origin divergence. The plots compare predictions of the evolutionary model on the extent of origin divergence (simulations of the *Lachancea* phylogenetic tree) with empirical data. **A:** Box plot of origins efficiency distributions split by family size. The plot compares origin families (sets of orthologous origins) in the nine *Lachancea* species (white line and red shaded areas) and in simulated species (blue boxes, for 100 simulation runs). Medians are shown as white line for data, black bar for simulation, 25-75 percentiles as shaded area for data, box for simulation, and 10-90 percentiles as coarse shaded area for data, whiskers for simulation. **B:** Origin divergence measured by the number of origins in the common ancestor that were lost in a pair of species, plotted as a function of total origin loss events. The plot compares model simulations (blue circles, 100 simulation runs), the experimental data (red squares) and a null model shuffling of empirical birth - death events in each branch (green triangles, 1000 simulation runs). Error bars are standard deviations on *x*−axis values. Simulation parameters (for the evolutionary model, see methods): *γ* = 2.3, *β* = 1.9, overall birth and death rate 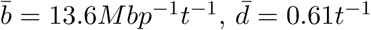 and rate of origin firing-rate reshuffling *R* = 0.92*t*^−1^, where *t* is measured by protein-sequence divergence.

As a second step, we have considered the model prediction for the divergence of the shared origins in two species descending from a common ancestor. Specifically, we asked whether the number of origin death events occurring in two branches of the tree could justify the number of common origins in the two species. Indeed, whenever in a pair of species the number of shared origins is lower than the number of origins belonging to their common ancestor, this discrepancy must be due to the evolutionary loss events. These events are predicted by our model to be correlated in diverging species, due to the common ancestry and the coupling of loss events to origin efficiency and distance. This correlation should lower the number of shared origins compared to a null expectation where loss events are not correlated. Fig. 4B shows that the model correctly predicts the experimental the divergence in the number of shred origins due to during evolution, without any parameter adjustment. We also verified that, as expected, a null evolutionary model is not able to reproduce this feature. The null model fixes in each branch of the simulated tree the same number of birth and death events that are present in the corresponding branch of *Lachancea* tree, but these events occur uniformly along the genome. The difference between the null model and the evolutionary model predictions shown in Fig. 4B is a consequence of correlated origins losses due to the common structure of sister species in terms of origins positions and efficiencies.

## DISCUSSION AND CONCLUSIONS

Overall, this study provides a framework to study replication-program evolution driven by replication-origin birth-death events, and demonstrates that both fork stalling and efficiency shape the adaptive evolution of replication programs. The model framework is predictive and falsifiable and it can be used to formulate predictions on the phylogenetic tree. In future studies, it would be interesting to explore the predictions for the evolutionary dynamics under perturbations, such as evolution under increased replication stress or conditions where fork stalling becomes more frequent. Additionally, the framework can be used to discover specific trends, such as different evolutionary dynamics of specific genomic regions (subtelomeres [23], regions containing repeats, etc. [24, 25]), role of genome spatial organization [26], and correlated firing of nearby origins.

The previous approach by Newman and coworkers [18] described the evolution of origin distance as an optimization process that minimizes double fork-stall events, without attempting to characterize explicitly the evolutionary dynamics. Such approaches are limited compared to the framework presented here, because they can predict only the origin-distance distribution, and they do not allow any prediction regarding origin and replication-program evolution along lineages and across phylogenetic trees. In accordance with the results of Newman *et al*., we confirm that double-stall events are a primary driver of the evolution of replication programs, and we frame this finding into the empirically measured birth-death evolutionary dynamics of replication origins. Additionally, we show that next to fork-stall events, origin efficiency plays an important role into shaping the evolutionary landscape seen by a replication timing profile.

What could be the mechanisms coupling efficiency to origin birth death? The actual process of origin death could be nearly neutral [27], as low-efficiency origins, are - by definition - rarely used, and unused origins, over evolutionary times are more prone to decay in sequence, and consequently in firing-rate until they disappear. Equally, a new-born origin close to a very strong one (and thus relatively inefficient) could be relatively less likely to establish and become stable over evolutionary times, compared to an isolated one, because it is used rarely. However rarely used origins could be essential in situations of stress (and in particular they could resolve double-stall events). Finally, a fitness cost for maintaining too many origins might set up an overall negative selection preventing a global increase in origin number [16, 28, 29].

## III. MATERIALS AND METHODS

### Data

The experimental data used in this work come from ref. [22]. In particular, we made use of the data regarding the replication origins. For each origin in each of the ten *Lachancea* species, this data set includes the chromosome coordinate and firing rate, and the inferred birth and death events occurred in the branches of the phylogenetic tree shown in Fig. S1. Focusing on the terminal branches of the tree and on the extant replication origins, this study defines three categories of origins: (i) “conserved” origins (which survived from the last ancestor) (ii) “newly gained” origins gained in the last branch of the phylogenetic tree, (iii) “lost” origins, which were present in the last ancestor species and are not present in the terminal branch. Properties of the lost origins (e.g. position and firing rate) are inferred from the projection of the corresponding ones on the closest species, keeping into account synteny. Since the synteny map is less precise in distant species, the information on the origins events is only available for the six sister species in the tree, which belong to the three closest species pairs, highlighted with the blue shaded area in Supplementary Fig. S1.

### Computation of the efficiency

Origin efficiency was defined as the probability of actively firing during S phase (or, equivalently, the probability of not being passively replicated by forks coming from nearby origins). In practice we computed it by the following formula

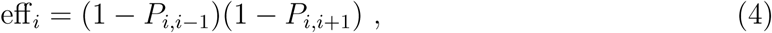

where *P*_*i,i*+1_ and*P*_*i,i*−1_ are the *i*-th origin probability of passive replication respectively from the (*i* + 1)-th and (*i* − 1)-th origin. Note that this efficiency formula Eq. 4, is an approximation that only takes into account the possibility to be passively replicated by neighbor origins, neglecting the influence of other nearby origins. Following ref [22], for computing the efficiency we assumed that the origin firing process has constant rate [16], and we thus obtain the following closed expressions for the probabilities of passive replication

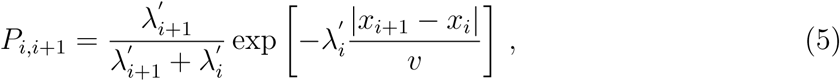

and

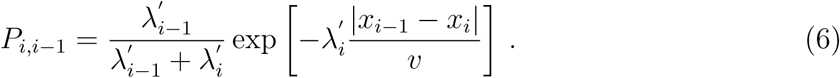

In the above equations, *v* is the typical velocity of replication forks, *x*_*i*_ is the *i*-th origin chromosome coordinate, and 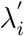 is the *i*-th origin firing rate divided by the mean firing rate of the species the origin belong to. The raw firing rates in the data are affected by the different physiology of the nine *Lachancea* species in the experimental growth conditions (which were the same for all the species). In order to reduce these differences, we normalized the rates by their average for each given species. For this reason, we did not make use of the origin efficiency data already present in [22].

### Computation of the double-stall probability

The probability 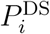 that two converging forks stall is easily computed in the limit where the stall probability per base-pair is small and the number of base-pairs is large. Under these assumptions, stalling is a Poisson process with rate (per base-pair) *π*. 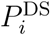 can be written in terms of the probability *P* ^S^(*x*) that a single fork stalls after replicating *x* nucleotides,

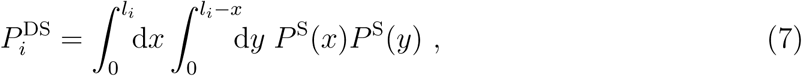

where *l*_*i*_ is the length (number of base-pairs) of the inter-origin region. Imagine two converging replication forks starting from origins *i* and *i* + 1: the two integration variables *x* and *y* represent the number of base-pairs that each fork replicates before stalling. By using the Poisson-process result *P* ^S^(*x*) = *π* exp(−*πx*) and performing the integration, one obtains the result in Eq. (1).

### Evolutionary model

We defined origin birth-death models incorporating different selective pressures. In these models, the genome is described as a one-dimensional circle with discrete origin location *x*_*i*_, where the length of the genome is equal to the average genome length in *Lachancea* clade (10.7*Mbp*). We made use of a circular genome in order to avoid border effects. In the model, the set of origins change over evolution by three basic (stochastic) processes, birth of an origin in a certain genome region, origin death and change of origins firing rate.

Overall origin birth/death rates were estimated from the data as follows. To estimate the overall birth rate 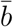 we considered, for all the terminal branches of the phylogenetic tree, the number of birth events *N*_*b*_, the genome length of the corresponding species *L* and the length of the tree branch *T*, and divided *N*_*b*_ by *LT*. Then we averaged over all terminal branches. To estimate the overall death rate 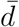, we followed a similar approach, taking the number of death events *N*_*d*_ in the terminal branches, the length of the branch *T* and the number of origins in the corresponding species *n*_*ori*_, then computing 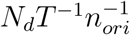 for all the terminal branches and averaging these values. The final results for overall birth and death rates from the origin birth death events across the *Lachancea* clade are 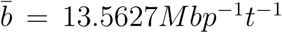 and 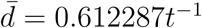.

The process by which origin firing rates change over evolution was described as stochastic, with every origin having a certain probability per unit time to change its firing rate, given by *R* = 0.92*t*^−1^, a value fitted from experimental data (see SI text and Fig. S3). When a firing rate changes, it is resampled from the distribution of all the empirical normalized firing rates, computed using the data in [22] (see SI text and Fig. S3 for more details).

### Simulations

#### Algorithm

The prediction of the different evolutionary models were derived numerically, making use of custom simulations written in C++, which implement the origin birth-death dynamics as a Gillespie algorithm [30]. The code is available with the authors. Every model variant was required to reproduce the experimental overall rates, 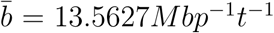 for origin birth, 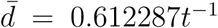 for origin death and *R* = 0.92*t*^−1^ for firing rate change. We simulated the three processes defining the model as follows. (i) The birth process has a common definition for the stall aversion and joint model. The algorithm first tests each subsequent inter-origin region, calculates the birth probability from Eq. 2 and stores the results. Subsequently, it computes the normalization factor *N*, in order to match the empirical birth rate per nucleotide 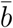. Finally, it samples all the inter-origin regions drawing birth events from the computed birth probability (Eq. 2). New origins are placed the mid points of the tested intervals. (ii) The death process is different for the stall-aversion model (unbiased) and the joint model (related to the origin efficiency). In the joint model, the algorithm first calculates the death rate for each origin using Eq. 3 and stores the results. Subsequently, it computes the normalization factor *N*, in order to match the empirical mean death rate 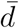. Finally, it samples all origin drawing death events from the computed death probability. For the unbiased process (stall-aversion model) the dynamics is identical, but all the origins have the same death rate 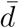, so that the algorithm can skip the calculation of *N*. (iii) The process updating origin firing rates over evolutionary times is common to all model variants. The probability of update per origin per unit time is *R*. Origins are sampled for each time step and assigned a new rate uniformly extracted from the empirical distribution of all normalized firing rates with probability *Rdt*.

During the simulation the genome configuration (chromosome position, firing rate, efficiency for each origin) is known at each time step, which matches the empirical time (tree-branch length, measured by protein-sequence divergence). For simulating single lineages, we started with a collection of 50 origins, with positions and firing rate uniformly drawn from all the possible ones. Rapidly, the number of origins, the inter-origin distances and the efficiency reach a steady state (set by the balance of birth and death rate, and characterized by approximately 225 origins). Configurations, including birth-death events, were printed at regular time intervals after steady state is reached. The time interval between prints is chosen to be equal to the average length of the *Lachancea* phylogenetic tree terminal branches, in order to compare single-lineage simulations with empirical data. For simulations on a phylogenetic tree, after one species reaches the steady state, it is used as a root. To reproduce the empirical branching structure of the tree, we run the simulation, one for each branch of the phylogenetic tree, each time starting from the species at the previous branching point, for a period that matches the length of the branch. If the simulated branch is terminal then the configuration corresponds to one of the empirical species, otherwise it corresponds to a “branching-point species” and it can be used as starting point for other simulations. Each simulation run gives nine different simulated species with the same cladogenetic structure as the empirical species (Fig. S1).

#### Fitting procedure

The biased birth-death processes in the simulations rely on some parameters to tune the strength of the bias, these are the only parameters to fix by a fit, since all the other parameter values are fixed empirically. In the joint model there are two free parameters, *β* and *γ* that tune respectively the strength of the bias on the origin birth and on the origin death process. For a discrete set of parameter pairs spanning realistic intervals we run hundred different simulations, each starting with a randomized genome. Considering the simulated species for all the pairs of parameter values, we quantify the discrepancy with experimental data by evaluating the L1 distance of the normalized histogram of efficiency and inter-origin distances. This quantity is a number between 0 and 2, 0 if the histograms perfectly overlap and 2 if they have completely different supports. For each pair of parameters the analysis gives two values of discrepancy. We choose the value of *γ* (the parameter that tunes the bias on the birth rate based on double-stall aversion) by taking the smaller discrepancy from the inter-origin distances distribution. For the value of *β* (which tunes the interference bias on the death rate in joint model), we chose the one that gave us the smaller area on the efficiency distribution. For the double-stall aversion model the fitting procedure is the identical, and only requires to fix *γ*.

### Null birth-death model

We defined a null birth-death model where origin birth-death events in sister species are uncorrelated, in order to analyze the divergence of shared origins and compare it with the prediction of the evolutionary model. This model implements birth and death events uniformly, regardless of origin position and firing rate, fixing the number of events for each branch of the simulated phylogenetic tree. These values are taken from the inference reported in ref. [22] (shown in Fig. 3A of that study and in Supplementary Fig. S1). The simulation of this model starts with 220 origins (the number of origins inferred for to LA2, the species at the root of the tree). Subsequently, following the structure of the *Lachancea* phylogenetic tree, the simulation proceeds as follows: (i) at each branching point the genome is copied into two daughters, (ii) for each daughter the prescribed number of random death and birth events (in this order) is generated on random origins (iii) the simulation stops when it reaches the leaves of the *Lachancea* tree.

## ACKNOWLEDGMENTS

We are very grateful to Ludovico Calabrese, Simone Pompei and Orso Maria Romano for the useful discussions. This work was supported by the Italian Association for Cancer Research, AIRC-IG (REF: 23258).

## Supplementary Figures

**FIG. S1.**
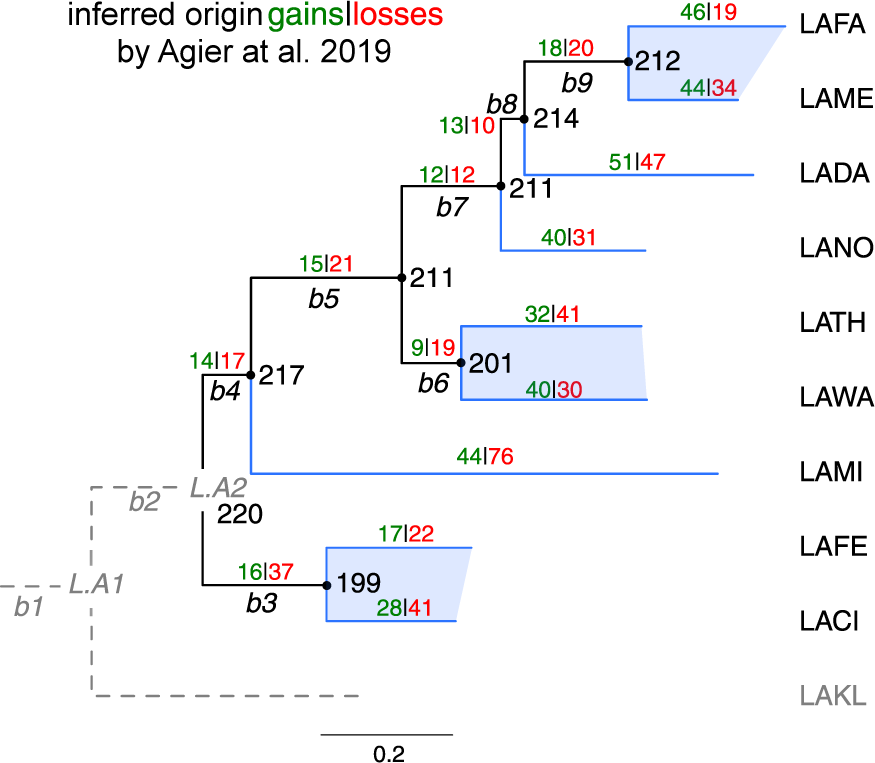
The phylogenetic tree of the ten *Lachancea* yeasts clade taken from ref [22], Fig.3A. the abbreviations of the species names are the following. LAFA: *L. fantastica*, LAME: *L. meyersii*, LADA: *L. dasiensis*, LANO: *L. nothofagi*, LATH: *L. thermotolerans*, LAWA: *L. waltii*, LAMI: *L. mirantina*, LAFE: *L. fermentati*, LACI: *L. cidri* and LAKL: *L. kluyveri*. LAKL was used as the outgroup species. Hence evolutionary events that occurred on both the LAKL and the b2 branches (dotted lines) could not be retraced. As a consequence, our simulations of the model set on the were not possible for the b2 and LAKL branches, and it was possible to simulate nine species instead of ten. Internal branches, labeled b3 to b9, and terminal branches are drawn in black and blue, respectively. The number of origin gains (in green) and losses (in red) were estimated for each branch of the tree in ref. [22]. The inferred number of active replication origins in the ancestral genomes is indicated next to the corresponding node of the tree. The six sister species, which belong to the three closest pairs of species, are highlighted with the blue shaded areas.

**FIG. S2.**
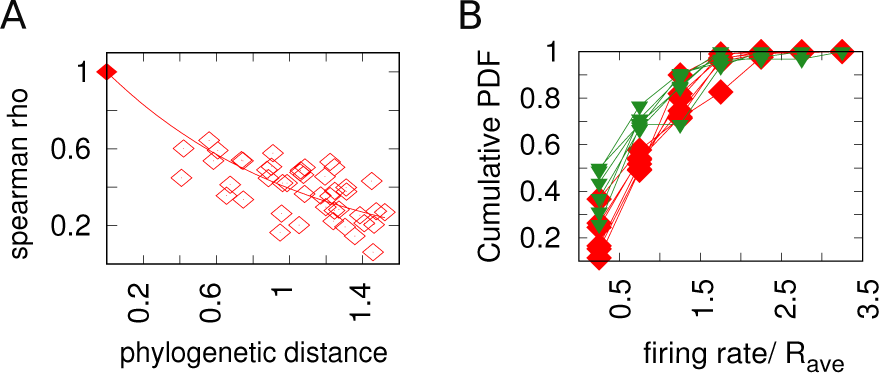
Experimental data on the evolutionary change of firing rates process. **A:** The firing rates Spearman correlation coefficient *ρ* between sets of corresponding origins decreases with increasing phylogenetic distance between species. Each point in the plot represents a pair of species. The *x* axis reports the phylogenetic distance between the two species, while the *y* axis reports the Spearman correlation between the sets of normalized firing rates for conserved origins between the two species. Empty squares represent the analysis carried out with *Lachancea* clade yeasts, while the symbol with coordinates (0,1) represents the fact that that non-distant species must have *ρ* = 1 **B:** Cumulative probability distribution of the normalized firing rates of newly-gained origins (green triangles) compared to the origins belonging to extant species (red squares), for the three sister species in the *Lachancea* clade. This plot shows that the two functions are very similar. This results is compatible with the assumption of resampling of firing rates over evolution taken for the model (see supplementary text).

**FIG. S3.**
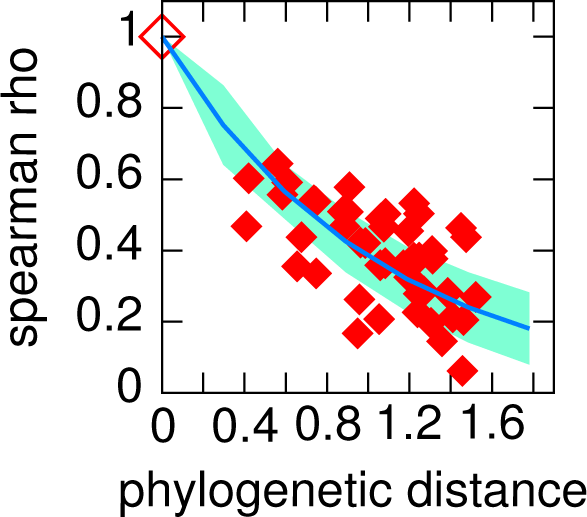
The decaying trend of the spearman correlation coefficient define a characteristic time for the firing rate resample. For each pair of species we compute the spearman correlation coefficient between the set of normalized firing rates belonging to corresponding origins. The figure shows the results of this analysis. The red points refer to experimental data, each dot is a pair of species, the x coordinate is the phylogenetic distance between them while the y one is the value of the spearman correlation coefficient. The squared empty dot in (0,1) is a fictitious point placed to remark that the spearman coefficient between non-distant species must be 1. The blue line represent the results of the simulation (1000 runs) with *R* = 0.92*t*^−1^ and the light blue area the standard deviation. We fixed the value of *R* by fitting this specific trend, and indeed the simulations that use this value of *R* show a remarkable agreement with the experimental trend. For the algorithm details see methods and supplementary text.

**FIG. S4.**
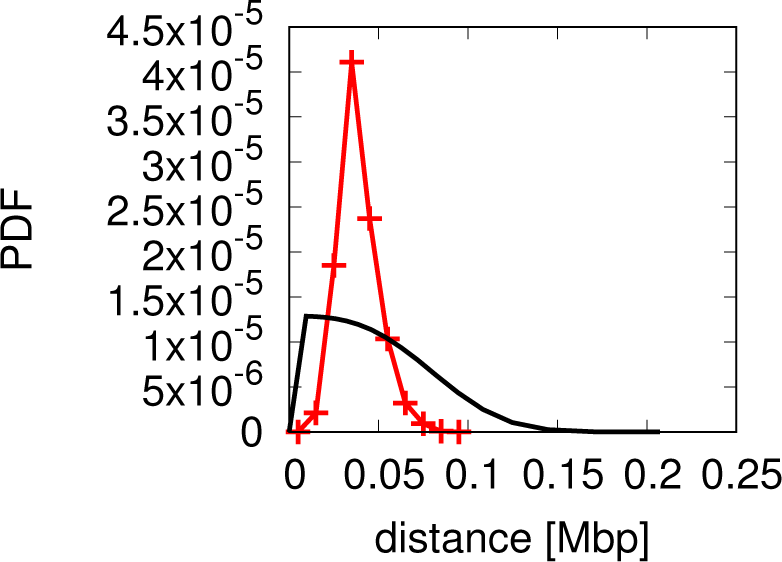
Analytical predictions for the inter-origins distance distribution falsify the scenario whereby interference alone drives replication-program evolution. The plot shows a comparison between the empirical inter-origin distance distribution (red line, crosses) and the analytical prediction from the scenario of origin birth-death driven by interference alone (black line, see methods and supplementary text for the calculation). The predicted distribution does not match the empirical one, thus the scenario can be rejected because it fails to reproduce a crucial feature of the data.

**FIG. S5.**
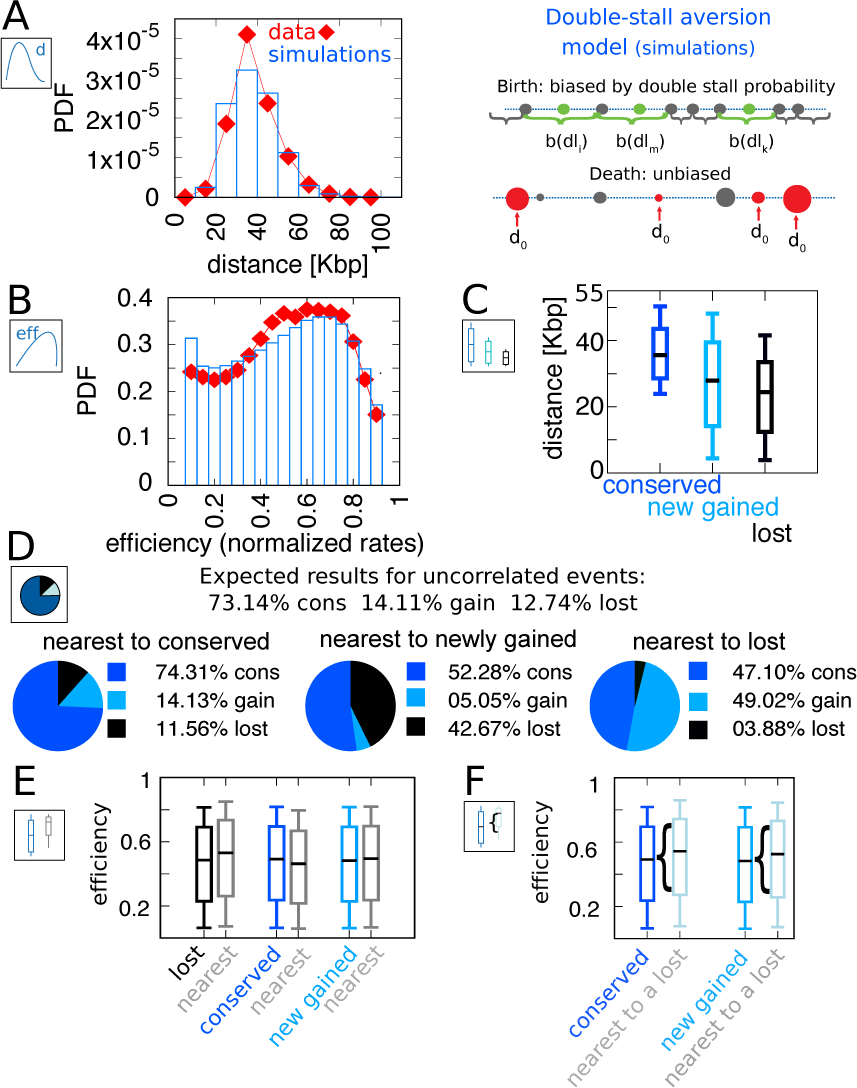
The double-stall aversion model reproduces origin turnover and distributions but fails to capture correlations between origin turnover and origin strength. The plots show the simulations of the best-fitting (*γ* = 2.4) model compared with empirical data. **A:** Inter-origin distance distribution in simulated species (blue bars) compared to the empirical distribution for the ten *Lachancea* species (red diamonds). **B:** Origin efficiency distribution in simulated (blue bars) *vs* empirical species (red diamonds). **C:** Box plot of the distance from the nearest origin split by evolutionary events, i.e. for conserved (dark blue), newly gained (blue) and lost origins (black), for simulated species. **D:** Fraction of origins that are nearest to conserved, newly gained and lost, for simulated species, compared to the expected result for uncorrelated events. **E:** Box plot of efficiency of lost, conserved and newly gained origins (respectively in black, dark blue and blue) and their neighbors (grey), in simulated species. The six distributions show very little variation. **F:** The efficiency of all conserved and newly gained origins compared to the ones flanking a lost origin. Box plots show the median (bar), 25-75 (box), and 10-90 (whiskers) percentiles. Simulation parameters (see methods): *γ* = 2.4, overall birth and death rate 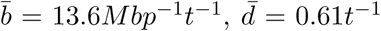 and firing-rate resampling rate *R* = 0.92*t*^−1^, where *t* is measured by protein-sequence divergence.

**FIG. S6.**
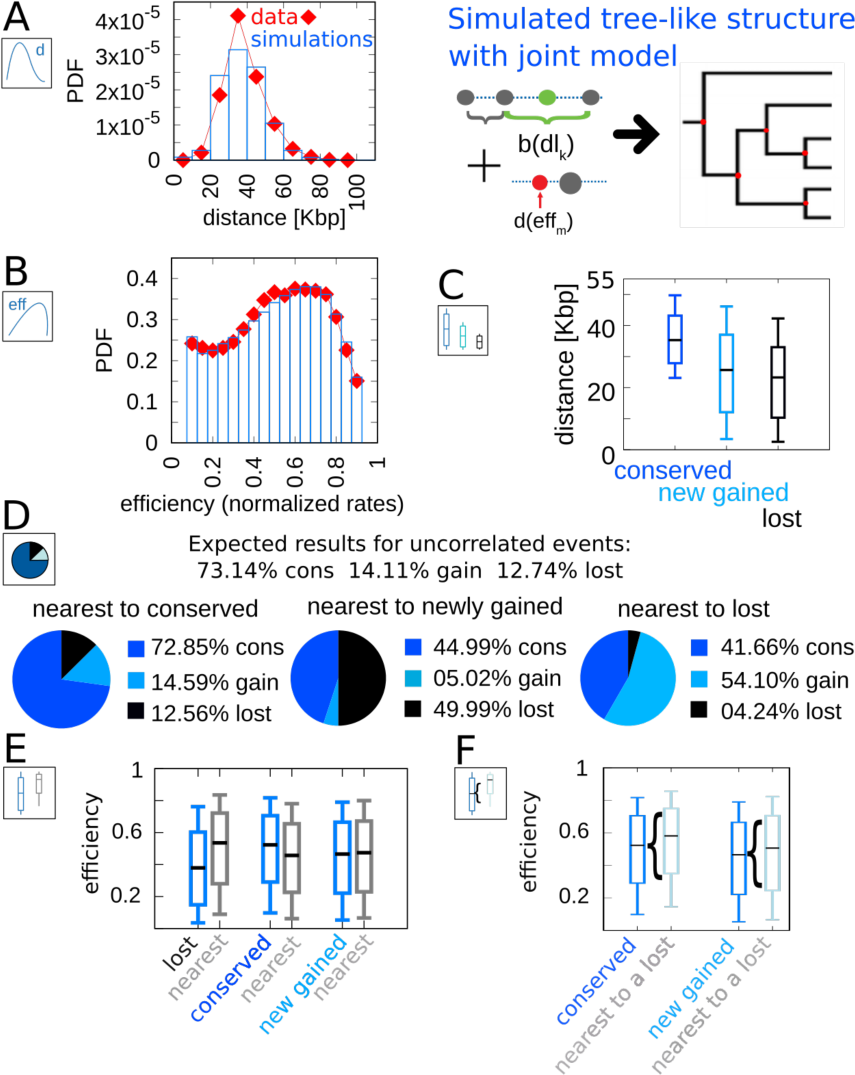
The joint efficiency/double-stall aversion model simulated on a cladogenetic structure reproduces all the results found for a single lineage. The results refer to 100 different runs of the simulation of the joint model on the empirical tree structure, compared with empirical data. **A:** Inter-origin distance distribution in simulated species (blue bars) compared to the empirical distribution for the ten *Lachancea* species (red diamonds). **B:** Origin efficiency distribution in simulated (blue bars) *vs* empirical species (red diamonds). **C:** Box plot of the distance from the nearest origin split by evolutionary events, i.e. for conserved (dark blue), newly gained (blue) and lost origins (black), for simulated species. **D:** Fraction of origins that are nearest to conserved, newly gained and lost, for simulated species, compared to the expected result for uncorrelated events. **E:** Box plot of efficiency of lost, conserved and newly gained origins (respectively black, dark blue and blue) and their neighbors (grey), in simulated species. **F:** The efficiency of all conserved and newly gained origins compared to the ones flanking a lost origin. Box plots show the median (bar), 25-75 (box), and 10-90 (whiskers) percentiles. Panels D and F show that the model correctly reproduces the correlation between origin birth-death events over evolution and efficiency of the nearest origin. Simulation parameters (see methods): *γ* = 2.3, *β* = 1.9, overall birth and death rate 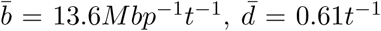 and rate of origin firing-rate reshuffling *R* = 0.92*t*^−1^, where *t* is measured by protein-sequence divergence.

## Supplementary Text

### A. Estimating parameters for the evolution of origin firing rates

This section motivates the model implementation of the evolutionary dynamics of firing rates. In order to quantify the change of origin firing rates over evolutionary times, we studied how the correlation between firing rates of conserved origins behave as species diverge (Fig. S2A). To quantify the divergence, for each pair of species in the *Lachancea* clade we calculated the Spearman correlation coefficient between the sets of firing rates belonging to corresponding origins in the two species considered (normalized by the species mean firing rate). We found that the more the species are distant, the less these two sets are correlated, which means that origin initiation rates diverge during evolution and origins loose memory of their initial firing rate. The model describes the evolution of firing rates as follows. Every origin changes its firing rate by extracting a new value from the distribution of empirical normalized ones, regardless of their previous firing rate. This process is characterize by a resampling rate *R*, common to all the origins, which defines the probability per unit time that an origin resamples its firing rate. The slope of the correlation coefficient in empirical data defines the speed at which the origin firing rates evolve. Hence, it is possible to fit this specific slope and extract the value of *R*.

In order to do that, we simulated the evolutionary process with unbiased origin death and update of the firing rate. This simulation can be performed without the birth process, because the only origins that one needs to consider in computing the Spearman coefficient between two species are the conserved ones. Each simulation started from 225 origins, with firing rates randomly sampled from the empirical set of firing rates, evolved the genomes changing the firing rates with the resampling process described above and removing the origins according to the death rate estimated from the data. By performing several simulations with different values of the extracting rate *R*, it is possible to fit its best value. For each *R* tested, we ran 1000 simulations for an evolutionary time corresponding to 1.6 times the maximal distance between species in the empirical *Lachancea* phylogenetic tree. After computing the Spearman correlations between snapshots at different evolutionary times, we performed an exponential fit, in order to see which value of the *R* parameter gave the best agreement with the experimental data, finding the best-fit value *R* = 0.92. Fig. S3 shows the trend achieved by the simulation using *R* = 0.92, and it shows a very good agreement between experimental data and simulations.

Note that in ref. [22], a similar analysis was carried out in order to verify if the reprogramming of the origins firing rate has an impact on the differentiation of replication timing. The authors analyzed the origin firing time *differences* between conserved replication origins in all pairs of species, and found that this difference does not correlate with the phylogenetic distance between species. This finding is apparently in contrast with our results, which suggest that origin reprogramming increases with distance between species. We believe that this discrepancy is due to the higher sensitivity of the Spearman correlation and of the use of species-average normalized firing rates in this study.

### B. The empirical data falsify the scenario where interference alone drives origin evolution

This section presents a theoretical analysis of the scenario where solely origin interference sets the evolutionary pressure on replication timing profiles. This analysis shows that a description that only takes into account the evolutionary pressure that acts on origin efficiency is not able to reproduce the origins spatial arrangement, a crucial feature in empirical yeast data. To carry out this analysis, we take a “maximum entropy” approach (see Banavar JR, Maritan A, Volkov I. Applications of the principle of maximum entropy: from physics to ecology. J Phys Condens Matter. 2010;22(6):063101. doi:10.1088/0953-8984/22/6/063101) and infer an effective “force potential” acting on inter-origin distance by looking at its (assumed equilibrium) distribution. Specifically, the effective potential acting on the origin efficiency starting from the empirical efficiency distribution, can be analytically computed from the following formula

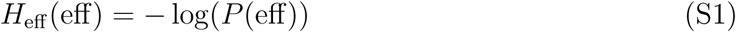

where eff is the efficiency, eff ∈ [0, 1], and *P* (eff) the efficiency probability density function.

The above potential, once given the relation between efficiency and distance between origins, Eq. 4, defines another potential *H*_*d*_(*d*) that act on the inter-origin distances. By taking the exponential of *H*_*d*_(*d*) one obtains the expected probability distribution predicted for the distances at equilibrium.

In order to find *H*_*d*_(*d*) one must to invert Eq. 4 and find *d*(eff). To accomplish this task, we have approximated the three-body interaction that gives the efficiency with a two-body interaction. This assumption implies that each origin feels the interference of only one of his two neighbors, and is effective as long as three-origin interactions can be decomposed in two-origin components. Under this assumption, Eq. 4 becomes

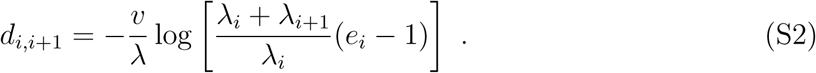

Note that origin efficiency, Eq. 4 also depends on the firing rates of the origin and its neighbor, hence, strictly speaking, one has that

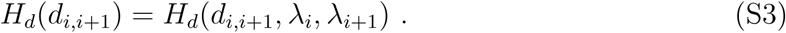

To eliminate the firing rates dependence we computed an effective potential 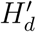 on the distance, which averages the effect of the different firing rates. To this end, we used the mean value theorem for integrals, as follows,

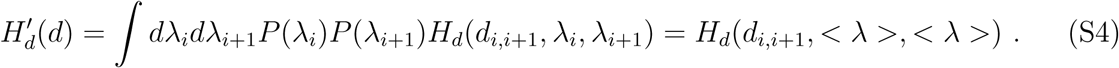

In other words we, substituted all the firing rates with the average one < *λ* >= 1, since the rates are normalized on the species average. With this simplification, going from *H*_eff_ (eff) to 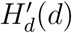 is straightforward, and gives

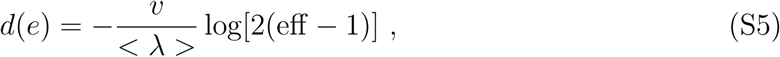

and

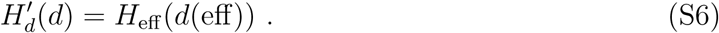

From the potential 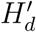, we can compute the prediction for the equilibrium probability distribution of inter-origin distances

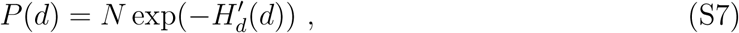

where *N* is a normalization factor. In order to use this calculation on the the data, we inferred the expected potential from the efficiency distribution, assuming that the interaction only depends on efficiency, and we then obtained the model prediction for the expected inter-origin distribution based on the efficiency profile. Comparison of this prediction with the empirical inter-origin distance distribution provides a test of the model. This procedure does not require to adjust any model parameter. Figure S4 shows the result of this analysis. The predicted distribution does not match the empirical one. This means that any evolutionary model that assumes a selective pressure based only on the efficiency (in other words, one that takes into account only the evolutionary pressure given by origin interference) cannot reproduce (at steady state) the correct spatial organization of replication origins.

